# Determining the interaction status and evolutionary fate of duplicated homomeric proteins

**DOI:** 10.1101/2020.04.08.031351

**Authors:** Saurav Mallik, Dan S Tawfik

## Abstract

Oligomeric proteins are central to life. Duplication and divergence of their genes is a key evolutionary driver, also because duplications can yield very different outcomes. Given a homomeric ancestor, duplication can yield two paralogs that form two distinct homomeric complexes, or a heteromeric complex comprising both paralogs. Alternatively, one paralog remains a homomer while the other acquires a new partner. However, so far, conflicting trends have been noted with respect to which fate dominates, primarily because different methods and criteria are being used to assign the interaction status of paralogs. Here, we systematically analyzed all *Saccharomyces cerevisiae* and *Escherichia coli* oligomeric complexes that include paralogous proteins. We found that the proportions of homo-hetero duplication fates strongly depend on a variety of factors, yet that nonetheless, rigorous filtering gives a consistent picture. In *E. coli* about 50%, of the paralogous pairs appear to have retained the ancestral homomeric interaction, whereas in *S. cerevisiae* only ∼10% retained a homomeric state. This difference was also observed when unique complexes were counted instead of paralogous gene pairs. We further show that this difference is accounted for by multiple cases of heteromeric yeast complexes that share common ancestry with homomeric bacterial complexes. Our analysis settles contradicting trends and conflicting previous analyses, and provides a systematic and rigorous pipeline for delineating the fate of duplicated oligomers in any organism for which protein-protein interaction data are available.

## Introduction

It is estimated that more than half of all proteins form oligomers. Oligomerization is thus ubiquitous and central to protein stability, function and regulation. Duplication is also ubiquitous and hence serves as the main source of new genes/proteins, as manifested by nearly half of all genes in a given genome being paralogs [1]. The duplication of genes encoding an oligomeric protein is of particular interest – the ancestral function may diverge alongside the oligomeric state thus providing new opportunities for evolutionary innovation [2–5].

Our analysis examined the divergence of homomers. By parsimony, the ancestors of both homomers and heteromers are homomers, as homomers are encoded by a single gene. Indeed, proteins have an inherent tendency to self-interact, and initially promiscuous self-interactions can be readily amplified by mutations to generate tightly bound homo-dimers and also larger homo-oligomers [6]. Upon duplication of a gene encoding a homomeric ancestor, and acquisition of the very first mutation(s), in either the original gene or its new copy, a statistical mixture of homo- and hetero-meric complexes would form [2] (**Fig 1A**, (***i***)>). Over time, further evolutionary divergence may result in three possible scenarios: (***ii***) loss of the capacity to cross-react and formation of two distinct homomeric complexes, or *obligatory homomers*; (***iii***) loss of the homomeric interactions and formation of a heteromeric complex, or *obligatory heteromers*. Alternatively, the interaction pattern may diverge asymmetrically – while one paralogue is kept as homomer, the other gains a completely new interaction partner (**Fig. 1A**, (***iv***): *hetero-others*). Other scenarios may occur, *e.g.*, loss of homomeric interactions in both copies, or divergence into monomers that do not interact with any other protein; however, these scenarios are intractable on a genome-scale (by parsimony, their ancestors cannot be assumed to be a homomer) and are probably relatively rare.

**Figure 1.**
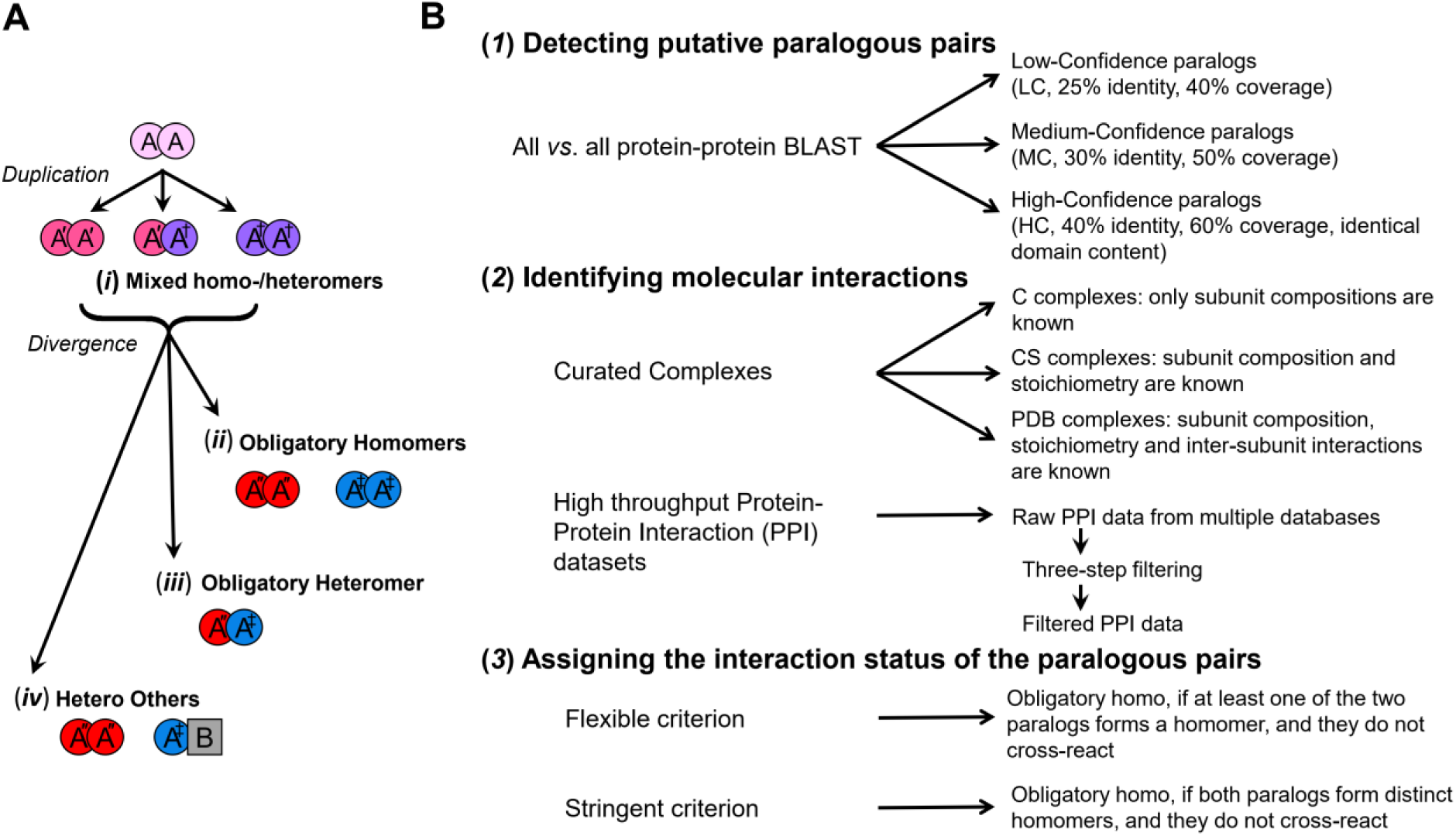
The potential evolutionary fates of duplicated homomeric proteins and the analysis pipeline for identifying them. **(A)** Duplication of a gene encoding a homomeric protein, and the emergence of the first mutation(s), leads to a statistical mixture of homo- and heteromeric complexes (*i*). Upon further divergence, three outcomes may arise: two distinct homomeric complexes (*ii*), a heteromeric complex involving both paralogs (*iii*), or loss of homomeric interaction in one copy, and gain of new interacting partners in the other paralog (*iv*). **(B)** Our analysis aimed to identify these four different evolutionary fates. It comprised three steps: (*1*) The genomes of *E. coli* and *S. cerevisiae* were each scanned to identify all possible paralogous protein pairs. These pairs were classified into three categories with increasing confidence of paralog assignment (note that all categories in our analysis are inclusive, *i.e.*, low-confidence paralogs include the medium-confidence ones, and the medium include the low-confidence pairs). (*2*) Interactions of these paralogs were identified and classified to homo- and heteromeric ones. Macromolecular complexes were collected from the Protein Data Bank (PDB complexes, inter-subunit interactions were obtained from crystal structure data) and the Complex Portal database (CS and C complexes, inter-subunit interactions were predicted from the PPI data). The *S. cerevisiae* PPI data were extracted from seven databases, and the *E. coli* data from eight databases. The raw PPI data were filtered using various criteria to exclude potential false-positives. *(3)* Finally, based on the identified interactions, the paralogous pairs were assigned to one of the four potential fates (*i*-*iv*, panel A) with either a *flexible* or a *stringent criterion*.

Individual cases following all these four scenarios are known. What remains unclear, however, is which fate is the most likely? Does protein function, or the source organism, for example, affect which fate dominates? Genome-scale studies [2–4] attempted to address the relative frequencies of these scenarios in model organisms, but their conclusions are inconsistent. Analyzing human, *Arabidopsis*, yeast and *E. coli* protein-protein interaction (PPI) data, [3] reported that most oligomeric paralogs diverged to form obligatory homomers. However, analysis of yeast, worm and fly, using both PPI data and oligomers of known structure, [2] indicated that heteromeric interactions dominate, a conclusion recently supported by [4] who analyzed yeast PPI data. We compared these studies and observed that these inconsistencies relate to three major factors. First, different evolutionary scenarios were examined in different studies – *e.g.*, ref.[3] did not consider the mixed homo/heteromers, and essentially none of these studies [2–4] consider hetero-others. Second, different interaction datasets were analyzed ranging from X-ray crystallographic structures (*e.g.*, ref.[2]) to high-throughput PPI data (*e.g.*, refs.[3,4]). Third, the divergence modes were assigned in an incongruous fashion. Following the definition of obligatory homomers, ref.[2] demanded both paralogs to be assigned as homomers; but others, refs.[3,4], for example, sufficed with identifying just one paralog as homomer.

Taking advantage of the extensive characterization of *Saccharomyces cerevisiae* and *Escherichia coli* macromolecular complexes, we investigated the potential evolutionary fates of their duplicated homomeric proteins. We systematically varied the stringencies of assigning paralogous pairs, of filtering molecular interaction datasets, and of assigning the divergence modes, and examined how these parameters affect the assigned proportions of homo-hetero divergence events. In *S. cerevisiae*, when stringent criteria were applied, a consistent picture arose, indicating that contrary to a previous analysis [3], 90% of duplications resulted in heteromeric complexes. In *E. coli*, however, it appears that paralogs are 5 times more likely to retain their ancestral homomeric interactions. We reconciled this difference by tracking down individual complexes and showing that complexes that are homomeric in *E. coli* have, upon duplication, diverged to heteromeric complexes in *S. cerevisiae*.

## Results

### A systematic approach to delineate the evolutionary fates of duplicated homomers

We analyzed the relative abundances of the four potential fates by examining the proteomes of *S. cerevisiae* and *E. coli* for which extensive interaction data exist. As the inconsistencies between previous works depict, this analysis presents biases at each one of its three steps (**Fig. 1B**). In the 1^st^ step, considering only paralogous pairs with high sequence coverage and identity would enrich closely related pairs that are more likely to be detected as mixed homo-heteromers. Conversely, assigning paralogous pairs with low coverage and identity might include cases where the changes in the divergence modes are due to loss or gain of entire domains rather than divergence of preexisting interfaces. To address this bias, in the 1^st^ step, we classified the putative paralogous pairs into three groups going from low to high confidence of paralogue assignment (**Fig. 1B, *1***).

In the 2^nd^ step, structures of macromolecular complexes allow to assign interactions with high accuracy, but crystal structures in particular create a bias in favor of homomeric interactions [7]. High-throughput protein-protein interaction (PPI) data cover a much larger set of proteins, yet they can be noisy, and how these data are filtered would substantially influence the results. Beyond random noise, there are biases – for example, certain PPI methods cannot detect homomeric interactions (*e.g.* pulldown and MS identification of binding partners). We thus analyzed separately and compared the results from curated complex datasets (hereafter, *curated complexes*) and high-throughput PPI data (**Fig. 1B, *2***). For the latter, PPI data were pulled together from different databases (**Data S1, S2**) and taken through three different filters to minimize false-positives. These databases encompass all reported interactions, including high resolution data, yet the high throughput data dominate, especially after the applied filtering.

Finally, in the 3^rd^ step, the criteria for assigning the fates of paralogous pairs also matter. In principle, obligatory-homomers means that *both* paralogs were individually observed as homomers and that a cross-interaction was not observed (*stringent criterion*). Sufficing with one paralog that forms a homomer would inevitably result in obligatory-homomers being the most frequent fate [4]. Further, as shown below, applying this *flexible criterion* results in assigning paralogs that actually diverged to hetero-others as obligatory-homo (**Fig. 1A, *iv*** and ***ii***, respectively). Thus, the divergence modes of the paralogous pairs were assigned applying both *stringent* and *flexible criteria* (**Fig. 1B, *3***). We subsequently examined the relative frequency of the four divergence modes, or fates, as a function of the stringency of analysis in each of the 3 steps.

Few clarifying notes regarding our analysis. We addressed paralogous pairs, *i.e.*, pairs of two genes that diverged from a common ancestor. In many cases, multiple paralogs exist that arose from two or more sequential duplications. Initially, we detected all potential pairs (**Fig. 1B**, step-1). Then, by assigning the divergence modes, we defined the relevant paralogous pairs (with few exceptions in the mixed category (**Fig. 1A, *i***) where one protein can be part of more than one pair). Thus, unless otherwise stated, the statistics and below discussion relate to gene pairs. Additionally, given that some complexes comprise multiple pairs, statistics are also provided per complexes. Finally, our parsimonious assumption is that the pre-duplicated ancestor can be considered a homomer if at least one descendent paralog is a homomer, and also if both paralogs are present as a heteromer (as in [2,4]). The latter was subsequently confirmed by our analysis (‘Yeast heteromeric paralogs diverged from bacterial homomeric ancestors’).

The results of our analysis were distilled to **Fig. 2** that presents the relative frequency of the four divergence modes given the dataset and stringency of analysis. The tables are arranged such that the darker the color, the higher is the stringency. The results given different stringencies of paralog assignment (Step-1) are presented in columns, going from low-confidence in pale green to high-confidence paralogs in dark green. Step-2, also in columns, from white (raw PPI data) to dark grey (Filter-3). Step-3, the stringency of assigning divergence modes, is presented in rows, with the top set of rows in yellow showing the *flexible criterion*, and the bottom, dark yellow rows indicating the *stringent criterion*. Finally, the dominant divergence modes, or fates, are highlighted in darker shades of red.

**Figure 2.**
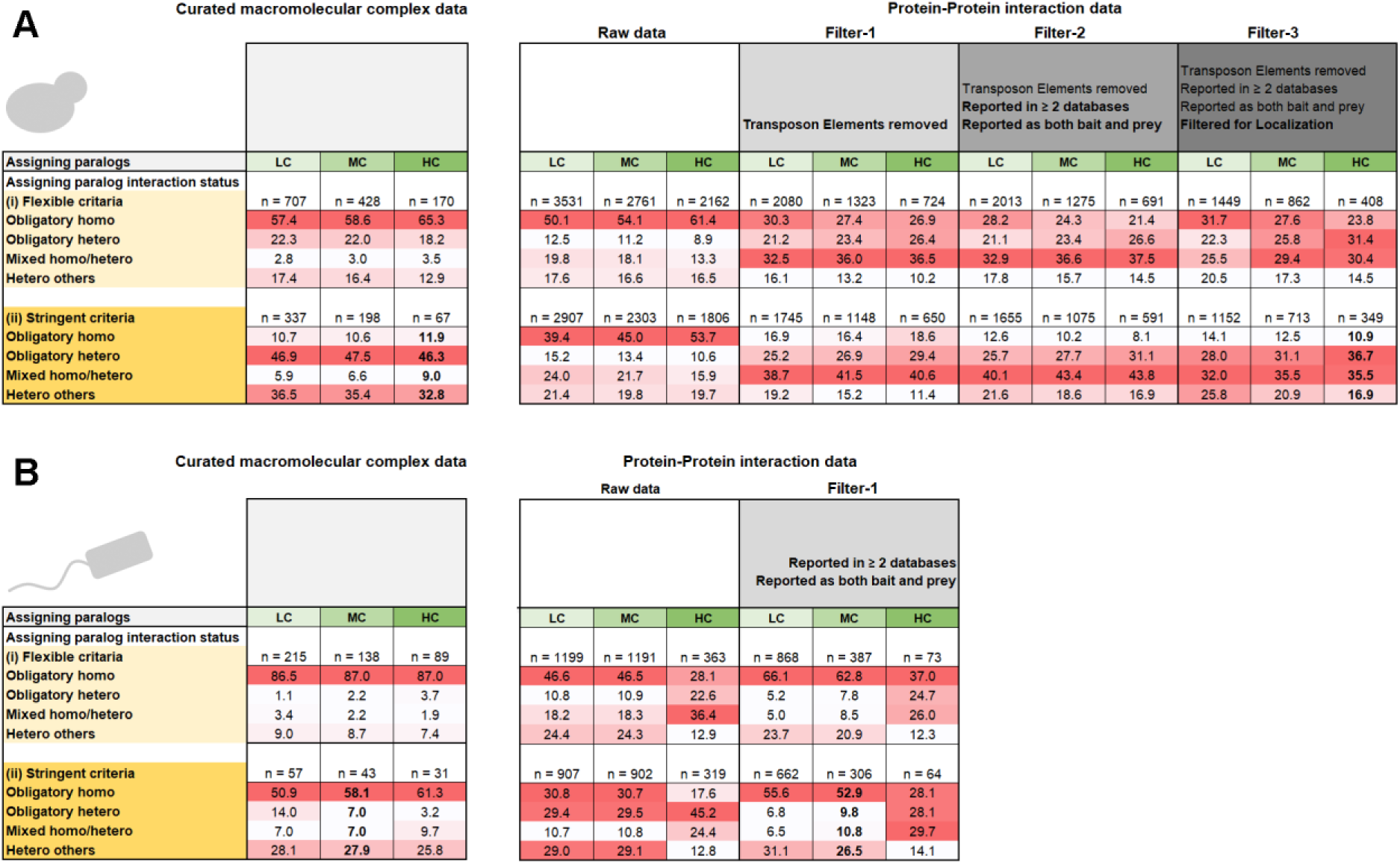
The distribution of divergence modes of *S. cerevisiae* and *E. coli* paralogous pairs. The four divergence modes, obligatory-homo, obligatory-hetero, mixed and hetero-others, are described in **Fig. 1A**. (**A**) The distribution of *S. cerevisiae* paralogous pairs in PPI data (*right panel*) and in curated complexes (*left panel*). Presented are the distributions for different stringencies of analysis, along its 3 steps (**Fig. 1B**). Step-1, paralog assignment, is presented in *columns*, shaded in green, from low-confidence in pale green to high-confidence paralogs in dark green. Step-2, identifying interactions, also in *columns*, from white (raw PPI data) to dark grey (filter-3). Step-3, the divergence mode, is presented in *rows* – the top set of rows represent the *flexible criterion* (shaded in yellow), and the bottom rows the *stringent criterion* (dark yellow). The dominant divergence modes, or fates, are highlighted in darker shades of red. (**B**) The distribution of *E. coli* paralogous.

### Heteromeric interactions dominate yeast paralogs

For yeast, under stringent filtering, the results from curated complexes and from PPI largely converge, indicating that ∼90% of yeast duplicates diverged to various heteromeric states. Specifically, stringent filtering of the PPI interactions (**Fig. 2A**, Filter-3, dark grey columns), and applying the *stringent criterion* for assigning the divergence modes (**Fig. 2A**, dark yellow rows), indicated that only about one-tenth of the paralogous pairs diverged to obligatory-homomers. Given the consistency between the two datasets, and the noise origins and biases indicated by our analysis (elaborated below), we surmise that obligatory-homo are indeed a minority in yeast (∼10%) and hetero-dominance is the reality (**Fig. 2A**, numbers in bold, **Data S3**). Within the three different hetero fates, the dominant fate is obligatory-hetero (about half of the pairs in the curated complexes, and a third in the PPI data where, as expected, a larger fraction of pairs was annotated as mixed).

If we were to count unique complexes instead of gene pairs, would the picture be different? Certain heteromeric complexes are composed of multiple paralogous proteins and these could shift the balance in favor of obligatory homomers (mostly ring-like complexes such as the proteasome; further addressed below). Nonetheless, analysis of complexes showed that, under the stringent filtering criteria, and for high-confidence paralogs, complexes comprising heteromers were nearly three-times more frequent than homomeric complexes (**Fig 3A**). Overall, we conclude that heteromeric interactions dominate yeast paralogs, regardless of whether we count paralogous pairs or unique complexes.

**Figure 3.**
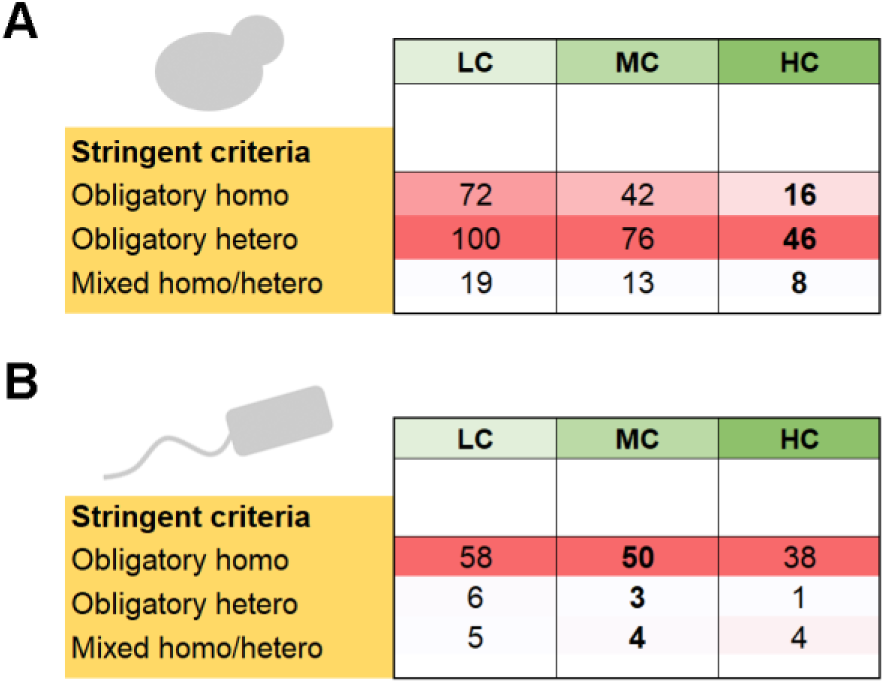
The distribution of complexes comprising homo- and heteromeric paralogs in *S. cerevisiae* and in *E. coli*. This analysis was based on the curated complexes databases. The column annotations and color shades are the same as in **Fig. 2**. (**A**) The numbers of unique *S. cerevisiae* complexes comprising paralogs assigned to the different homo/hetero divergence modes. Note that the different confidence levels for paralog assignment (LC, MC, HC) show that same trend as in **Fig. 2B**, curated complex panel. (**B**) The same for *E. coli*.

### Data biases and their mitigation

Our analysis also reveals various sources of error and bias, and how these could be mitigated. As expected, consistency of the two interaction datasets, curated complexes and PPI, fades away at lower stringency. Foremost, the 3^rd^ step of the analysis, assigning the divergence modes, had a massive impact on the relative abundances of homo-hetero pairs. Assigning obligatory homomers using the *flexible criterion* (suffice that one paralog is a homomer and no cross-reaction) resulted in ∼5-fold proliferation of obligatory-homomers in the curated complexes, and ∼3-fold proliferation in the PPI data (**Fig. 2A**, light yellow rows). The reason being that under the *flexible criterion*, hetero-others were assigned as obligatory-homo. Thus, cases that are quite abundant in yeast where one paralog kept the ancestral homomeric interaction and the other diverged to bind a completely new partner were not only ignored, but also mis-assigned.

Our analysis also reflects the homo- or hetero-biases that are inherent to the source of interaction data. The homo-dominance in the curated complexes dataset primarily stems from the known bias of crystal structures to detect homomers [7]; the hetero-dominance in the PPI dataset stems from certain high-throughput methods failing to detect homomeric interactions. Indeed, for a given a stringency with respect to the first two steps of the analysis (assigning paralogs, identifying interactions), homomers are more frequent in the curated complexes while heteromers dominate the PPI data (**Fig 2A**). However, these biases seem to be alleviated under the *stringent criterion*, as both the PPI and the curated complexes give a similar distribution of fates. Thus, consideration of all four evolutionary fates, namely including both mixed homo-hetero and hetero-others, is critical, as are adequate criteria to assign them (re the *stringent criteria*).

Two other elements seem to be critical for obtaining consistent results, both relating to the PPI data. Upon manual inspection we noticed that five long terminal repeat retrotransposon families, comprising a total of 90 proteins. These paralogous mobile genetic elements of viral origin [8,9] caused an inflation in the fraction of obligatory-homomers (∼50%, that dropped to ∼15% once removed). Further, once these retrotransposon proteins were removed (**Fig. 2A**, filter-1), the homo-hetero fates in the PPI data converged with those in the curated complexes (**Fig. 2A**, stringent criterion). Filtering of potential false-positives in the PPI data had a lesser effect. First, we applied a demand that interactions are reported in two different databases, and that interactions were detected with the protein pairs applied as both bait and prey (**Fig. 2A**, filter-2). The second source of false-positives are *in vitro* PPI interactions that do not occur *in vivo*. Obvious cases include interactions between proteins localized in different compartments (**Fig. 2A**, filter-3). However, compared to the removal of retrotransposons these two filters had a minor effect.

Overall, we conclude that heteromeric interactions between paralogous pairs is the dominant fate in yeast, regardless of whether we count paralogous pairs or unique complexes.

### Homomeric interactions dominate *E. coli* paralogs

A similar analysis of *E. coli* indicated that in oppose to *S. cerevisiae*, for high-confidence paralogs, about 60% of the descendent pairs are obligatory-homers in the curated complexes compared to only 30% in the PPI data (**Fig 2B**, filter-2, HC, *stringent criterion*, **Data S4**). However, this inconsistency is because the *E. coli* sample sizes for high-confidence paralogs are too small (**Fig. S1A**). In yeast, filtering led to considerable reduction in sample sizes, yet these remained high even for high-confidence paralogs (**Fig. 2A**). Further, the distribution is similar for high and medium-confidence, and with few exceptions even to the low-confidence (highest sample size, **Fig. S1B**). This is not the case for the *E. coli* analysis. When more distantly related paralogs were removed (MC and HC columns), sample sizes decreased by >10-fold, compared to >3-fold in yeast. Indeed, in yeast, owing to the relatively recent whole genome duplication, high-confidence paralogs comprise ∼60% of all detectable paralogs (1806/2907, **Fig. S1B, C**), while in *E. coli* they comprise only ∼35% (**Fig. S1B**). Thus, it seems that medium-confidence paralogs better report the actual reality in *E. coli*.

Overall, considering the stringent criterion for assigning the divergence fates, the filtered PPI data and the curated complexes gave a consistent picture by which ∼55% of the pairs comprise obligatory-homomers, for both medium- and low-confidence paralogs (**Fig 2B**, MC and LC). Further, as in yeast, homomers also dominated when complexes were counted (**Fig 3B**). Overall, it appears that retaining the ancestral homomeric interaction is the most likely fate of *E. coli* gene duplications.

Note that tuning the stringencies in the *E. coli* analysis had similar effects as in *S. cerevisiae*. Filtering the PPI for interactions reported in at least two databases, and as both bait and prey resulted in a higher fraction of obligatory-homomers. On the other hand, assigning the divergence modes with a *flexible criterion* resulted in overestimation of obligatory-homomers (and a corresponding drop in obligatory-heteromers).

### Yeast heteromeric paralogs diverged from bacterial homomeric ancestors

We observed the dominance of obligatory-homomers in *E. coli* (∼50%) while in *S. cerevisiae* they comprise only ∼10% of the duplicated oligomeric proteins, and in turn obligatory-heteromers comprise the majority. However, these two model organisms share common ancestry, as reflected in about one-third of *S. cerevisiae* proteins, many of which are mitochondrial proteins, harboring sequence signatures of bacterial origin [10]. We thus searched for the *E. coli* orthologs of the *S. cerevisiae* heteromeric paralogs, asking which are homomeric.

A systematic reciprocal BLAST was performed between all known *E. coli* homomers (n = 1033) and all *S. cerevisiae* obligatory-hetero and mixed paralogous pairs (n = 692; out of a total of 1152 LC pairs in the *stringent* categories, PPI dataset; **Fig. 2A**). Following manual curation (see **Methods**), we identified about a third of the heteromeric yeast paralogous pairs that have *E. coli* homomeric orthologs (n = 235; **Data S5**). Of these, nearly two-thirds, 153 pairs, relate to *E. coli* homomers that are singletons (*i.e.*, non-duplicated genes; a total of 52 proteins). By parsimony, these reflect cases of duplication and divergence of an ancestral bacterial homomer into paralogous heteromers in yeast. Remarkably, 42/52 of these *E. coli* proteins are metabolic enzymes that duplicated and diverged into heteromeric *S. cerevisiae* enzymes. In many such cases only one copy retained the catalytic activity whereas the other one evolved into a regulatory subunit. Examples include mitochondrial NAD^+^-dependent isocitrate dehydrogenase complex [11], Trehalose Synthase Complex [12], the 20S proteasome core particle subunits [13] or the ATP-dependent 6-phospho-fructokinase complex [14,15]. Other enzymes, such as chaperonins, HSP70 chaperones, and DNA and RNA helicases appear to have gone through multiple duplications and contribute to the hetero-dominance in *S. cerevisiae*.

The remaining third, 82 yeast heteromeric paralogous pairs, are orthologous to 144 obligatory homomeric pairs in *E. coli* (**Data S5**). These also relate to divergence of homomers to heteromers. What is unclear though is which of these genes duplicated independently in these two clades, and which one diverged to heteromers in an earlier bacterial ancestor. What is clear though is that the dominance of heteromeric paralogs in yeast is the result of homomers duplicating and preferentially diverging into heteromers.

## Discussion

With the obvious caveat of being based on two model organisms for which extensive protein interactions data are available, our analysis indeed suggests a continuous evolutionary process of bacterial homomeric proteins gradually duplicating and diverging into heteromeric proteins in eukaryotes. This ongoing evolutionary transition also validates our assignment of the fundamentally different divergence modes of paralogous pairs in *E. coli* and *S. cerevisiae* (**Fig. 2**). Assuming *E. coli* and *S. cerevisiae* are representatives of bacteria and single-cell eukaryotes, the gene duplications that occurred in the eukaryotic lineage that diverged from bacteria via endosymbiosis [16,17] led to 5-fold decrease in the abundance of homomers among paralogous proteins. Further, because paralogous proteins comprise nearly half of the proteomes, this phenomenon has led to a complete shift from the prevalence of homomers in prokaryotes to heteromers in eukaryotes [18,19].

The transition of homomeric prokaryotic complexes into eukaryotic heteromeric ones was previously noted for individual protein families, and especially for ring-like complexes such as DNA/RNA helicases [20,21], TCP complex subunits [22], proteasome [23,24] and exosome [25]. However, examining our dataset revealed that both ring-like and non-ring-like prokaryotic homomers evolved into heteromeric complexes in eukaryotes, and by a single or multiple gene duplications (**Fig. 4**, **Data S5**). Thus, the dominance of heteromeric paralogs in *S. cerevisiae* is not only because the ancestral homomers duplicated and diverged into heteromers, but also because heteromeric paralogs further duplicated and their descendants retained the heteromeric state.

**Fig 4.**
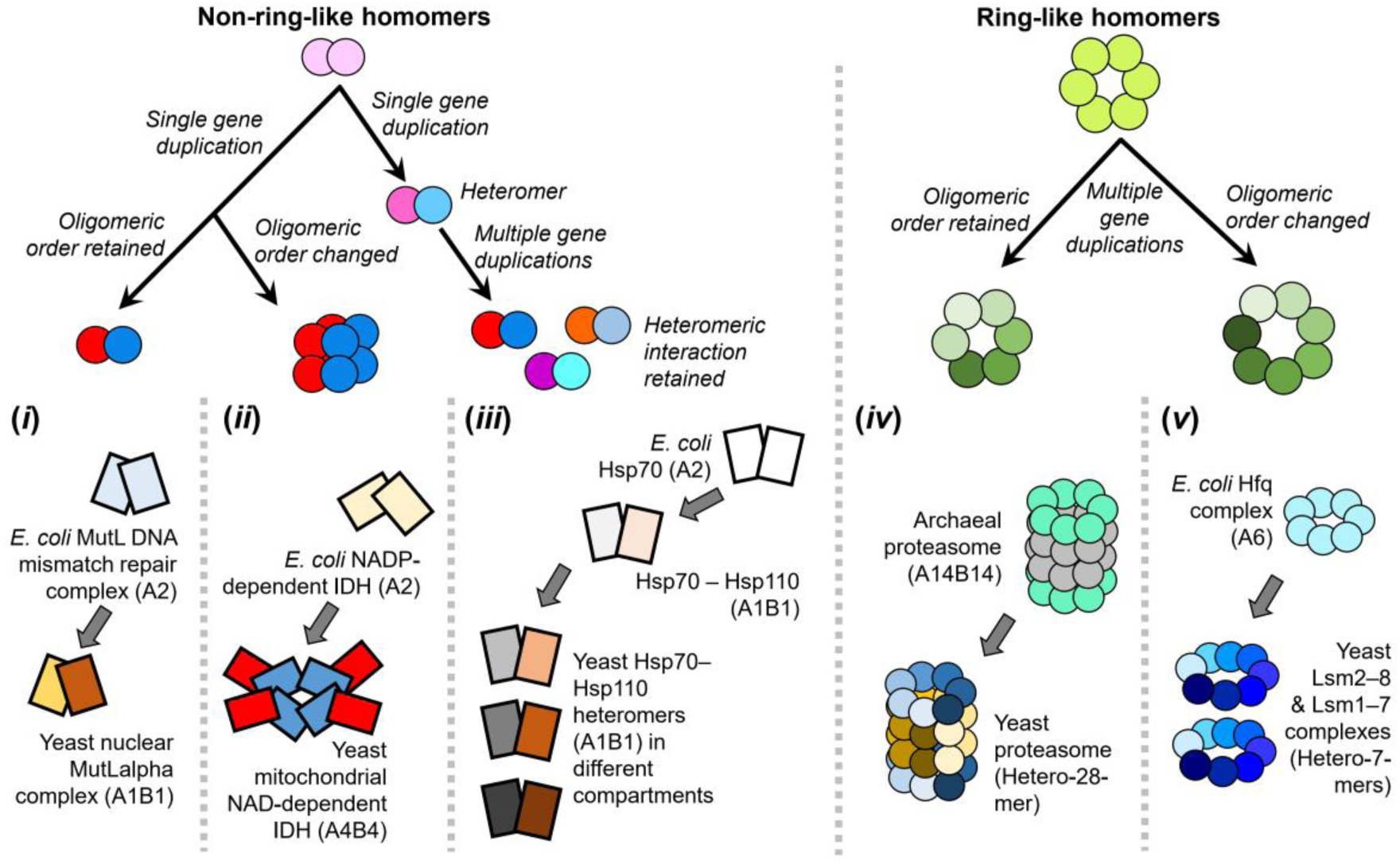
Different modes of prokaryotic homomer to eukaryotic heteromer transition. Gene duplication of an ancestral non-ring homomer may produce a heteromeric complex that may (***i***) or may not (***ii***) retain the ancestral oligomeric order (*i.e.*, the total number of subunits in the complex). After the first gene duplication and the subsequent emergence of a heteromeric interaction, multiple rounds of duplication may follow in which the descendant paralogs retain the heteromeric interaction (***iii***). For ring-like complexes, multiple rounds of intra-ring gene duplications result in heteromeric rings, while keeping (***iv***) or changing the ancestral oligomeric order (***v***). For each mode of transition, an example case is provided.

For non-ring-like complexes, a single gene duplication typically results in a single eukaryotic heteromeric complex that may or may not retain the ancestral oligomeric order (total number of complex subunits). For example, *E. coli* DNA mismatch repair endonuclease MutL is a homodimer, and the yeast orthologue is a heterodimer [26] (**Fig. 4, *i***). On the other hand, the bacterial homo-dimeric isocitrate dehydrogenase [11] duplicated and diverged into a hetero-octameric mitochondrial isocitrate dehydrogenase in yeast [26] – namely, the oligomeric order changed from 2 to 8 (**Fig. 4, *ii***). In this case, duplication and divergence into a heteromer tendered the opportunity of evolving a new regulatory mode by diversifying one subunit, while the other subunit kept the catalytic activity.

As a prokaryotic non-ring-like homomer evolves into a heteromer in eukaryotes, multiple rounds of duplication may occur and the descendent paralogs retain the newly evolved heteromeric interaction (**Fig. 4, *iii***). For example, the bacterial homomeric Hsp70 that duplicated and diverged into Hsp110 co-chaperones in eukaryotes [27,28], and the *S. cerevisiae* genome encodes multiple copies of Hsp70 and Hsp110 that form distinct heteromers in different subcellular compartments [29].

For ring-like prokaryotic homomeric complexes (e.g., helicase, protease, RNase and chaperonins), homo-to-hetero transition predominantly also occurred while retaining the ancestral oligomeric order or modifying it (**Fig. 4, *iv-v***). Complexes that have retained their ancestral oligomeric order (**Fig. 4, *iv***) include the archaeal homo-hexameric MCM complex that became hetero-hexameric in eukaryotes [21], and the core proteasome alpha- and beta-rings that remained heptameric [23,24]. In contrast, the bacterial helicase homo-hexameric Hfq ring-complex [30] diverged to the hetero-heptameric Lsm1-7 and Lsm2-8 complexes in yeast (**Fig. 4, *v***).

The above-described phenomena that underline homo-to-hetero transitions present some interesting questions. This transition needs to overcome the inherent self-interacting tendency of proteins, and eventually lead to incompatibility of the homomeric interactions. It is therefore likely to be adaptive, *i.e.*, provide a distinct functional advantage [6]. In *E. coli* duplications primarily yield obligatory homomers, with each paralog mediating a different enzymatic function (typically different substrate specificity). In yeast, however, the obligatory heteromers seem to be associated with acquisition of new regulatory modes. Thus, function may dictate the fate of the oligomeric state. Another factor might be the location of the active-site that in some enzymes resides within the subunits and in others at the interface between subunits [31]. Also of note is that, in principle, divergence of a heteromeric interaction increases the likelihood that both copies would fix in the genome, because loss of one copy leads to non-functionalization. Duplication itself is random, yet whether a duplicate is fixed or lost (the far more likely fate) depends on how rapidly it provides a selectable advantage [32]. Gene knockout experiments support this hypothesis – deletion of one copy is highly deleterious in heteromers while for obligatory homomers deletion of one copy often has little effect (**Data S3**).

Future work might address the above and other questions, and may also track down other possible evolutionary transitions – *e.g.* the dominating trend is homo-to-hetero transitions, yet can we track down cases of heteromers that diverged to homomers? Addressing these questions will demand detailed phylogenies and experimental evaluation of the oligomeric states before and after the duplication. However, a rigorous way of assigning oligomeric states from molecular interaction databases, and of determining the fate of duplicates, is crucial to any such investigation.

## Methods

Further details are provided in the supplementary items, in relation to the each of analyses described therein.

### Detecting *S. cerevisiae* and *E. coli* paralogous protein pairs

The 1^st^ step of our analysis identified all *S. cerevisiae* and *E. coli* paralogous protein pairs (**Fig. 1B**). To this end, all-versus-all intra-species protein-protein BLAST [33] was performed across their respective proteomes, obtained from NCBI Genome Database [34]. BLAST hits associated with at least 25% identity and 40% query coverage were manually inspected and assigned as putative paralogous pairs (3958 pairs in *S. cerevisiae* and 2090 pairs in *E. coli*, **Fig. S1**). These pairs were further classified into three overlapping groups, with increasing stringency of paralogue assignment, Low-Confidence (LC, ≥25% identity, ≥40% coverage), Medium-Confidence (MC, ≥30% identity, ≥50% coverage) and High-Confidence (HC, ≥40% identity, ≥60% coverage, and identical domain content). To ensure identical domain content, we compared the Pfam [35]–annotated domain contents of all HC pairs. Pfam uses Hidden Markov Models to identify domains and every annotated instance is given a probability score (*p*-value). Any domain assigned with *p* < 10^−5^ significance was considered for further analysis. Following domain assignments, paralogous pairs were compared and those that differ in their domain content were discarded. The list of 455 *S. cerevisiae* ohnologs (paralogs emerging from the whole genome duplication; **Fig. S1**) were collected from the Yeast Gene Order Browser [36].

### Identifying molecular interactions

#### Curated complexes

Curated homo- and hetero-meric macromolecular complexes of both *S. cerevisiae* and *E. coli* were collected from Protein Data Bank [37], 3D complex database [38] and Complex Portal [26]. Complexes that include at least one protein annotated as paralog were classified into three groups, with increasing stringency of curation accuracy (**Data S1, S2**). The first group, C complexes, comprises 127 *S. cerevisiae* and 18 *E. coli* complexes annotated in Complex Portal, for which only the subunit composition data are available (subunit stoichiometry is either unknown or only partially known). The second group, CS complexes, comprises 83 *S. cerevisiae* and 33 *E. coli* complexes annotated in Complex Portal, for which both subunit composition and stoichiometry data are available. The third group, PDB complexes, includes 167 *S. cerevisiae* and 117 *E. coli* complexes collected from the Protein Data Bank, for which subunit composition, stoichiometry as well as interaction patterns are known. The subunit stoichiometry of PDB complexes were further cross-validated by 3D complex database [38] annotations.

#### Protein-protein interaction data

For *S. cerevisiae*, 721701 binary PPI data were collected from seven different databases: BioGRID [39], DIP [40], HiNT [41], IntAct [42], iRefIndex [43], Mentha [44], and STRING [45] and 123644 interactions involving paralogous pairs were extracted. For *E. coli* 47727 binary PPI data was compiled, by adding one additional dataset [46] to the above seven databases; 4376 interactions involve paralogous proteins. Note that these PPI databases include both high- and low-throughput data, with the former dominating (see also next section). Predicted interactions, and text-mining based interactions, reported in STRING were removed. *S. cerevisiae* raw PPI data were filtered in three successive steps (**Data S1**). In the 1^st^ step, 90 transposon element proteins encoded by genes of viral origin that are included in the Saccharomyces Genome Database [47] were removed. In the 2^nd^ step, to minimize false-positives in the PPI data, we demanded that the interaction between two proteins observed using both proteins as bait and as prey, and the interaction must be reported in at least two databases. The bait-and- prey information is relevant to high-throughput two-hybrid and pull down experiments, and hence this filtering criterion removed interactions detected by other means, foremost by low-throughput methods such as gel shifts. However, this filtering resulted in a negligible loss of interacting pairs and did not bias the results (see next section). Also note that the databases used here collect their raw data from published literature. Overlaps between databases are therefore common, although none of these databases overlap completely. Thus, the demand that the interaction must be reported in at least two databases does not necessarily mean two independent observations, but as a minimum it eliminates annotation mistakes. In the 3^rd^ filtering step, interactions between two proteins localized in different sub-cellular compartments were excluded. For this, yeast protein localization data obtained from LoQAtE [48], Yeast GFP Fusion Localization Database [49] and Yeast Protein Localization database [50] were combined together. These filtering steps resulted in the final PPI dataset of 28381 pairwise interactions involving paralogous proteins (**Data S1**). The PPI datasets derived after each step of filtering are provided in **Data S1**. Transposon elements were absent in the *E. coli* raw PPI data and filtering involved only one step (interactions must be reported for both proteins as bait and as prey, and in at least two databases). This yielded a final PPI dataset of 1996 pairwise interactions (**Data S2**).

### Assigning the interaction status of paralogous pairs

Based on the interactions in the above-described molecular interaction datasets, paralogous pairs were assigned to one of the four categories described in **Fig. 1A**: obligatory hetero (the two paralogs do not self-interact, but cross-react to form a heteromer), mixed homo/hetero (two paralogs cross-react to form a heteromer, and at least one paralog also self-interacts), or hetero others (only one paralog self-interacts and the other interacts with to another, non-paralogous partner). Obligatory homomers were assigned using a *stringent* and a *flexible criterion*. The *stringent criterion* demanded that the two paralogs do not cross-react, and that *both* self-interact; the *flexible criterion* demanded that the two paralogs do not cross-react and at least one of them self-interacts.

PDB structures and PPI data, by definition, comprise physical interaction data between proteins. For CS and C complexes, inter-subunit interactions were predicted from the PPI data. A homomer was assigned if it is present in multiple copies in a complex, and also self-interacts in the PPI data. Heteromers were assigned if both paralogs co-occur in a complex and found to cross-interact, but not self-interact, in the filtered PPI dataset. For obligatory homomers in the curated complexes, we also ensured that the two paralogs do not cross-interact in the PPI data.

To examine if the assignment of homo/hetero fates in the PPI dataset was substantially influenced by the bait-prey filtering, we extracted all the PPI data that were detected exclusively by methods other than two-hybrid and pull downs. As with other PPI data, these data were filtered by demanding that the interaction is reported in at least two databases, and by excluding interactions between proteins localized in different sub-cellular compartments. For the filtered subset of data, applying the *stringent criterion*, we assigned the interaction status of paralogous pairs. Only 14 new pairs were detected (**Data S6**), compared to 1152 pairs in the filtered PPI data (see **Data S3**), indicating that a negligible amount of PPI data was lost due to the bait-prey filtering. Further, in these 14 pairs, the overall dominance of heteromers in yeast was reflected (5 obligatory hetero, 7 mixed, 2 hetero-others, and no obligatory homo; **Data S6**).

### *S. cerevisiae* and *E. coli* orthologous proteins

To identify the orthologous *S. cerevisiae* and *E. coli* protein pairs, inter-species reciprocal protein-protein BLAST [33] searches were performed. In total, 7325 protein pairs associated with e-value < 10^−5^ were extracted. We then identified the subset of these pairs that comprise a homomeric protein in *E. coli* and an obligatory-hetero, or mixed homo-hetero paralogous protein in *S. cerevisiae*. The domain content of these pairs, as annotated in Pfam [35], were compared and those sharing at least one domain, were extracted. These pairs were then manually checked for having the same function in the two organisms and that the shared domain corresponds to this function. When consolidated, this analysis extracted orthologous relationships between 103 *E. coli* homomeric proteins (52 singletons and 144 paralogous pairs) and 421 paralogous *S. cerevisiae* proteins (235 pairs; **Data S5**).

### Statistical analysis

All the computation and statistical analyses were performed using in-house Python codes. Graph plots were generated using OriginLab software and Adobe Photoshop.

## Acknowledgements

We sincerely thank Emmanuel Levy and Christian Landry for critically reading of an early version of this manuscript and for their valuable comments and suggestions. This work is supported by the Estate of Mark Scher. S.M. was supported by PBC-VATAT Postdoctoral Fellowship, provided by the Council for Higher Education, Israel.

## Author Contributions

S.M. and D.S.T. designed research; S.M. implemented computational methodologies and analyzed data; S.M. and D.S.T. discussed and commented on the results and wrote the manuscript.

## Data availability

All data associated with this work are provided as Supplementary Information (Data S1–S5, Fig. S1).

**Correspondence and requests for materials** should be addressed to D.S.T.

## Supplementary Figure legends

**Fig. S1. Histogram plots of *E. coli* and *S. cerevisiae* paralogous pairs binned by sequence identity. (A)** *E. coli* paralogs (n = 2090 pairs). **(B)** *S. cerevisiae* all paralogs (n = 3958 pairs). **(C)** *S. cerevisiae* ohnologs (the subset of paralogs that arose from the whole genome duplication; n = 455 pairs). Note that these plots include all paralogs, not only the ones for which molecular interaction data are available. The dotted red lines represent the identity thresholds used for defining MC (≥30% identity) and HC (≥40% identity).

## Supplementary data file legends

**Data S1.** The *S. cerevisiae* molecular interaction dataset used in this study (including the list of the curated complexes and the PPI data).

**Data S2.** The *E. coli* molecular interaction dataset used in this study (including the list of the curated complexes and the PPI data).

**Data S3.** The inferred interaction status of *S. cerevisiae* paralogous pairs, in curated complexes and in PPI data. For paralogous pairs in curated complexes, deletion phenotypes are also provided.

**Data S4.** The inferred interaction status of *E. coli* paralogous pairs, in curated complexes and in PPI data.

**Data S5.** List of S. cerevisiae heteromeric paralogs that relate to homomeric E. coli proteins.

**Data S6.** The inferred interaction status of *S. cerevisiae* paralogous pairs in PPI data detected exclusively by methods other than two-hybrid and pulldowns.

## References

1. Penel S, Arigon AM, Dufayard JF, Sertier AS, Daubin V, Duret L, et al. Databases of homologous gene families for comparative genomics. BMC Bioinformatics. 2009;10: 1–13. doi: 10.1186/1471-2105-10-S6-S3

2. Pereira-Leal JB, Levy ED, Kamp C, Teichmann SA. Evolution of protein complexes by duplication of homomeric interactions. Genome Biol. 2007;8. doi: 10.1186/gb-2007-8-4-r51

3. Hochberg GKA, Shepherd DA, Marklund EG, Santhanagoplan I, Degiacomi MT, Laganowsky A, et al. Structural principles that enable oligomeric small heat-shock protein paralogs to evolve distinct functions. Science (80-). 2018;359: 930–935. doi: 10.1126/science.aam7229

4. Marchant A, Cisneros AF, Dube AK, Gagnon-Arsenault I, Ascendo D, Jain H, et al. The role of structural pleiotropy and regulatory evolution in the retention of heteromers of paralogs. Elife. 2019;8: 1–34. doi: 10.7554/eLife.46754

5. Diss G, Gagnon-Arsenault I, Dion-Coté AM, Vignaud H, Ascencio DI, Berger CM, et al. Gene duplication can impart fragility, not robustness, in the yeast protein interaction network. Science (80-). 2017;355: 630–634. doi: 10.1126/science.aai7685

6. Lukatsky DB, Shakhnovich BE, Mintseris J, Shakhnovich EI. Structural Similarity Enhances Interaction Propensity of Proteins. J Mol Biol. 2007;365: 1596–1606. doi: 10.1016/j.jmb.2006.11.020

7. Marsh JA, Teichmann SA. Structure, Dynamics, Assembly, and Evolution of Protein Complexes. Annu Rev Biochem. 2015;84: 551–575. doi: 10.1146/annurev-biochem-060614-034142

8. Carr M, Bensasson D, Bergman CM. Evolutionary Genomics of Transposable Elements in Saccharomyces cerevisiae. PLoS One. 2012;7. doi: 10.1371/journal.pone.0050978

9. Bourque G, Burns KH, Gehring M, Gorbunova V, Seluanov A, Hammell M, et al. Ten things you should know about transposable elements. Genome Biol. 2018;19: 199. doi: 10.1186/s13059-018-1577-z

10. Ku C, Nelson-Sathi S, Roettger M, Sousa FL, Lockhart PJ, Bryant D, et al. Endosymbiotic origin and differential loss of eukaryotic genes. Nature. 2015;524: 427–432. doi: 10.1038/nature14963

11. Wang P, Lv C, Zhu G. Novel Type II and Monomeric NAD+ Specific Isocitrate Dehydrogenases: Phylogenetic Affinity, Enzymatic Characterization, and Evolutionary Implication. Sci Rep. 2015;5: 1–11. doi: 10.1038/srep09150

12. Bell W, Sunt W, Hohmann S, Wera S, Reinders A, De Virgilio C, et al. Composition and functional analysis of the Saccharomyces cerevisiae trehalose synthase complex. J Biol Chem. 1998;273: 33311–33319. doi: 10.1074/jbc.273.50.33311

13. Bochtler M, Ditzel L, Groll M, Hartmann C, Huber R. The proteasome. Annu Rev Biophys Biomol Struct. 1999;28: 295–317. doi: 10.1146/annurev.biophys.28.1.295

14. Poorman RA, Randolph A, Kemp RG, Heinrikson RL. Evolution of phosphofructokinase - Gene duplication and creation of new effector sites. Nature. 1984;309: 467–469. doi: 10.1038/309467a0

15. Banaszak K, Mechin I, Obmolova G, Oldham M, Chang SH, Ruiz T, et al. The crystal structures of eukaryotic phosphofructokinases from Baker’s yeast and rabbit skeletal muscle. J Mol Biol. 2011;407: 284–297. doi: 10.1016/j.jmb.2011.01.019

16. Eme L, Spang A, Lombard J, Stairs CW, Ettema TJG. Archaea and the origin of eukaryotes. Nat Rev Microbiol. 2017;15: 711–723. doi: 10.1038/nrmicro.2017.133

17. Zaremba-Niedzwiedzka K, Caceres EF, Saw JH, Bäckström Di, Juzokaite L, Vancaester E, et al. Asgard archaea illuminate the origin of eukaryotic cellular complexity. Nature. 2017;541: 353–358. doi: 10.1038/nature21031

18. Marsh JA, Teichmann SA. Protein Flexibility Facilitates Quaternary Structure Assembly and Evolution. PLoS Biol. 2014;12. doi: 10.1371/journal.pbio.1001870

19. Bergendahl LT, Marsh JA. Functional determinants of protein assembly into homomeric complexes. Sci Rep. 2017;7: 1–10. doi: 10.1038/s41598-017-05084-8

20. Veretnik S, Wills C, Youkharibache P, Valas RE, Bourne PE. Sm/Lsm genes provide a glimpse into the early evolution of the spliceosome. PLoS Comput Biol. 2009;5: 1–14. doi: 10.1371/journal.pcbi.1000315

21. Bochman ML, Schwacha A. The Mcm Complex: Unwinding the Mechanism of a Replicative Helicase. Microbiol Mol Biol Rev. 2009;73: 652–683. doi: 10.1128/mmbr.00019-09

22. Gupta RS. Evolution of the chaperonin families (HSP60, HSP 10 and TCP-1) of proteins and the origin of eukaryotic cells. Mol Microbiol. 1995;15: 1–11. doi: 10.1111/j.1365-2958.1995.tb02216.x

23. Gille C, Goede A, Schlöetelburg C, Preißner R, Kloetzel PM, Göbel UB, et al. A comprehensive view on proteasomal sequences: Implications for the evolution of the proteasome. J Mol Biol. 2003;326: 1437–1448. doi: 10.1016/S0022-2836(02)01470-5

24. Wollenberg K, Swaffield JC. Evolution of proteasomal ATPases. Mol Biol Evol. 2001;18: 962–974. doi: 10.1093/oxfordjournals.molbev.a003897

25. Lykke-Andersen S, Brodersen DE, Jensen TH. Origins and activities of the eukaryotic exosome. J Cell Sci. 2009;122: 1487–1494. doi: 10.1242/jcs.047399

26. Meldal BHM, Bye-A-Jee H, Gajdoš L, Hammerová Z, Horácková A, Melicher F, et al. Complex Portal 2018: Extended content and enhanced visualization tools for macromolecular complexes. Nucleic Acids Res. 2019;47: D550–D558. doi: 10.1093/nar/gky1001

27. Easton DP, Kaneko Y, Subjeck JR. The Hsp110 and Grp170 stress proteins: Newly recognized relatives of the Hsp70s. Cell Stress Chaperones. 2000;5: 276–290. doi: 10.1379/1466-1268(2000)005<0276:THAGSP>2.0.CO;2

28. Dragovic Z, Broadley SA, Shomura Y, Bracher A, Hartl FU. Molecular chaperones of the Hsp110 family act as nucleotide exchange factors of Hsp70s. EMBO J. 2006;25: 2519–2528. doi: 10.1038/sj.emboj.7601138

29. Bogumil D, Alvarez-Ponce D, Landan G, McInerney JO, Dagan T. Integration of two ancestral chaperone systems into One: The evolution of eukaryotic molecular chaperones in light of eukaryogenesis. Mol Biol Evol. 2014;31: 410–418. doi: 10.1093/molbev/mst212

30. Sauter C, Basquin J, Suck D. Sm-like proteins in Eubacteria: The crystal structure of the Hfq protein from Escherichia coli. Nucleic Acids Res. 2003;31: 4091–4098. doi: 10.1093/nar/gkg480

31. Abrusán G, Marsh JA, Wilke C. Ligand-Binding-Site Structure Shapes Allosteric Signal Transduction and the Evolution of Allostery in Protein Complexes. Mol Biol Evol. 2019;36: 1711–1727. doi: 10.1093/molbev/msz093

32. Finnigan GC, Hanson-Smith V, Stevens TH, Thornton JW. Evolution of increased complexity in a molecular machine. Nature. 2012;481: 360–364. doi: 10.1038/nature10724

33. Altschu SF, Gish W, Miller W, Myers EW, Lipman DJ. Basic Local Alignment Search Tool. J Mol Biol. 1990;215: 403–410.

34. Benson DA, Cavanaugh M, Clark K, Karsch-Mizrachi I, Lipman DJ, Ostell J, et al. GenBank. Nucleic Acids Res. 2013;41: 36–42. doi: 10.1093/nar/gks1195

35. Finn RD, Bateman A, Clements J, Coggill P, Eberhardt RY, Eddy SR, et al. Pfam: The protein families database. Nucleic Acids Res. 2014;42: 222–230. doi: 10.1093/nar/gkt1223

36. Byrne KP, Wolfe KH. The Yeast Gene Order Browser: Combining curated homology and syntenic context reveals gene fate in polyploid species. Genome Res. 2005;15: 1456–1461. doi: 10.1101/gr.3672305

37. Berman H, Henrick K, Nakamura H, Markley JL. The worldwide Protein Data Bank (wwPDB): Ensuring a single, uniform archive of PDB data. Nucleic Acids Res. 2007;35: 2006–2008. doi: 10.1093/nar/gkl971

38. Levy ED, Pereira-Leal JB, Chothia C, Teichmann SA. 3D complex: A structural classification of protein complexes. PLoS Comput Biol. 2006;2: 1395–1406. doi: 10.1371/journal.pcbi.0020155

39. Oughtred R, Stark C, Breitkreutz BJ, Rust J, Boucher L, Chang C, et al. The BioGRID interaction database: 2019 update. Nucleic Acids Res. 2019;47: D529–D541. doi: 10.1093/nar/gky1079

40. Salwinski L. The Database of Interacting Proteins: 2004 update. Nucleic Acids Res. 2004;32: 449D – 451. doi: 10.1093/nar/gkh086

41. Das J, Yu H. HINT: High-quality protein interactomes and their applications in understanding human disease. BMC Syst Biol. 2012;6. doi: 10.1186/1752-0509-6-92

42. Orchard S, Ammari M, Aranda B, Breuza L, Briganti L, Broackes-Carter F, et al. The MIntAct project - IntAct as a common curation platform for 11 molecular interaction databases. Nucleic Acids Res. 2014;42: 358–363. doi: 10.1093/nar/gkt1115

43. Razick S, Magklaras G, Donaldson IM. iRefIndex: A consolidated protein interaction database with provenance. BMC Bioinformatics. 2008;9: 1–19. doi: 10.1186/1471-2105-9-405

44. Calderone A, Castagnoli L, Cesareni G. Mentha: A resource for browsing integrated protein-interaction networks. Nat Methods. 2013;10: 690–691. doi: 10.1038/nmeth.2561

45. Szklarczyk D, Morris JH, Cook H, Kuhn M, Wyder S, Simonovic M, et al. The STRING database in 2017: Quality-controlled protein-protein association networks, made broadly accessible. Nucleic Acids Res. 2017;45: D362–D368. doi: 10.1093/nar/gkw937

46. Rajagopala S V., Sikorski P, Kumar A, Mosca R, Vlasblom J, Arnold R, et al. The binary protein-protein interaction landscape of escherichia coli. Nat Biotechnol. 2014;32: 285–290. doi: 10.1038/nbt.2831

47. Cherry JM, Hong EL, Amundsen C, Balakrishnan R, Binkley G, Chan ET, et al. Saccharomyces Genome Database: The genomics resource of budding yeast. Nucleic Acids Res. 2012;40: 700–705. doi: 10.1093/nar/gkr1029

48. Breker M, Gymrek M, Moldavski O, Schuldiner M. LoQAtE - Localization and Quantitation ATlas of the yeast proteomE. A new tool for multiparametric dissection of single-protein behavior in response to biological perturbations in yeast. Nucleic Acids Res. 2014;42: 726–730. doi: 10.1093/nar/gkt933

49. Huh WK, Falvo J V., Gerke LC, Carroll AS, Howson RW, Weissman JS, et al. Global analysis of protein localization in budding yeast. Nature. 2003;425: 686–691. doi: 10.1038/nature02026

50. Kals M, Natter K, Thallinger GG, Trajanoski Z, Kohlwein SD. YPL.db2: The Yeast Protein Localization database, version 2.0. Yeast. 2005;22: 213–218. doi: 10.1002/yea.1204

