## Supplementary material for "Determining the interaction status and evolutionary fate of duplicated homomeric proteins": Fig. S1

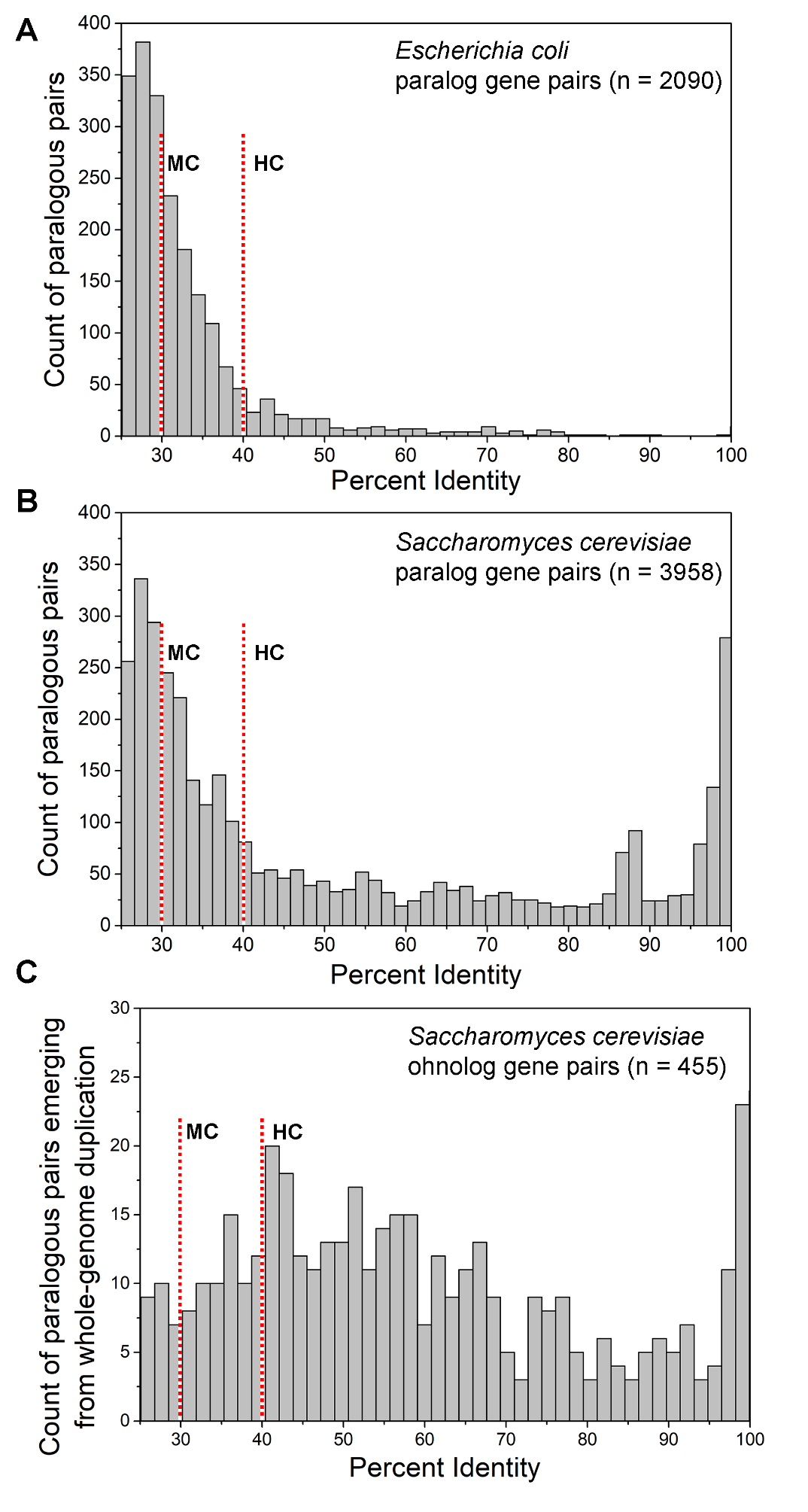

**Fig. S1**. **Histogram plots of E. coli and S. cerevisiae paralogous pairs binned by sequence identity. (A)** *E. coli* paralogs (n = 2090 pairs). **(B)** *S. cerevisiae* all paralogs (n = 3958 pairs). **(C)** *S. cerevisiae* ohnologs (n = 455 pairs), *i.e.*, the subset of paralogs that emerged from whole genome duplication. Note that these plots include all paralogs, not only the ones for which molecular interaction data are available. The dotted red lines represent the identity thresholds used for defining MC (≥30% identity) and HC (≥40% identity).
